# Quantitative light element (sodium and potassium) profiling in plant tissues using monochromatic X-ray fluorescence analysis

**DOI:** 10.1101/2024.10.22.619597

**Authors:** Parvinderdeep S. Kahlon, Pinelopi Kokkinopoulou, Silvia Bugallo Alfageme, Cheuk Ka Leong, Zewu Chen, Christa Testerink, Antony van der Ent

## Abstract

Accurately determining the elemental composition of plant tissues is essential for physiological studies on plant stress, including salinity tolerance. However, high-throughput routine analysis of light elements (range of sodium to calcium) in plant samples is challenging due to the need for complete sample dissolution and expensive inductively coupled plasma-mass-spectrometry (ICP-MS) analysis. Lower costs method (ion chromatography, ion selective electrodes) exists, but also require sample dissolution and lack sensitivity for very small samples (<10 mg). This study reports on a new method for the quantitative analysis of light elements in plant tissues using monochromatic X-ray fluorescence (XRF) instrumentation and innovative sample preparation and mounting. We used this approach to assess elemental uptake, distribution, and accumulation in *Arabidopsis thaliana* and *Oryza sativa* plants subjected to salt stress. The method can be used on samples as small as 1 mg making it suitable for small *Arabidopsis thaliana* plants. We systematically evaluated different sample preparations methods, repeatability, and measurement times to confirm the robustness of the technique. The results show that the monochromatic XRF method delivers rapid, non-destructive, and extraction-free analysis, strongly correlating with ICP-MS acquired data. As such, the monochromatic XRF method is a reliable and efficient alternative for studying salinity tolerance ideally suited for investigating elemental composition of early plant developmental stages, offering new possibilities for research into early stimuli sensing, perception and nutrient efficiency.

## INTRODUCTION

Plants are essential for sustaining life on Earth. The nutrients they absorb directly determine their ability to grow, thrive, and support other life forms. In plant nutrition, a range of elements can be defined as either essential or beneficial to a varying degree (Brown et al., 2022). Out of the 17 essential elements for plant growth, 11 can be categorized light elements, defined here as the elements sodium (*Z* = 11) to calcium (*Z* = 20). These light elements play a crucial role throughout the life cycle of plants in growth and development, especially nitrogen (N), phosphorus (P) and potassium (K), which are the most common components in commercial fertilizers. Addition of these elements in terrestrial ecosystem have shown to be beneficial for above ground biomass (Fang et al., 2024). However, in saline soils, the abundance of elements such as sodium (Na) and chloride (Cl) are increased, and their uptake by plants leads to toxicity due to competition with beneficial elements (Zhao et al., 2020). Depending on the level of salinity in soil, this exchange of beneficial to disruptive ion uptake, and the decrease in water uptake can lead to severe consequences from defective phenotypes in plants to whole crop loss (Zörb et al., 2018).

Plants uptake these elements in form of ions and this uptake is regulated by the ion transport. This transport is either through specific transporters or passive diffusion to maintain cellular balance. Under saline conditions, Na^+^ competes with essential ions such as K^+^ and Ca^2+^, creating an imbalance in physiological functions. To maintain cellular function, plants use mechanisms like Na^+^/K^+^ antiporters to exclude Na or transport it into vacuoles, minimizing its toxic effects (Barzana et al., 2021; Van Zelm et al., 2020). Significant efforts have been invested in understanding the mechanisms and role of these transporters in enhancing salt tolerance, a critical trait for improving plant performance, a key characteristic importance for agricultural production (Zhao et al., 2020). In various crops at different developmental stages, quantitative trait loci (QTL) related to salinity stress have been identified, particularly involving Na⁺ and K⁺ transport genes (Afzal et al., 2023). Overall, the mechanisms of salt tolerance involve a combination of exclusion of Na⁺ from the cytosol, maintenance of K⁺ uptake, and regulation of osmotic potential (Munns & Tester, 2008). Consequently, one effective approach involves analysing elemental concentrations, such as Na and K, across diverse germplasm to assess the potential for developing new salt-tolerant breeding lines.

To perform elemental analysis, various methods has been developed to date. Conventional methods such as inductively coupled plasma-mass-spectrometry (ICP-MS), ion chromatography (IC) and ion selective electrodes (ISE), which have their individual advantages and limitations (Rieger and Litvin, 1998; Goyal et al., 1993). ICP-MS offers the highest sensitivity with low limits of detections, but necessitates acid digestion of the plant material prior to analysis (Masson et al., 2010). This is time-consuming, expensive, and requires a digestion microwave (Wilschefski and Baxter et al., 2019; Djingova et al., 2013). The use of IC and ISE also requires time-consuming sample preparation involving complete solubilisation of the plant material, pH adjustment and ionic strength adjustment (for ISE) (Frenzel and Michalski, 2016). None of the conventional methods are particularly high throughput and suffer from incomplete dissolution of the sample matrix. This is especially important for light elements, notably silicon (Si) and aluminium (Al) unless hydrofluoric acid digestion is used, which is a highly hazardous laboratory procedure (Taber et al., 2002).

X-ray fluorescence spectroscopy (XRF) does not require solubilization of plant material, and avoiding digestion makes it potentially high throughout (Hossain et al., 2021). In combination with simple sample grinding, a handheld XRF instrument can achieve an accuracy comparable to digestion-based methods with inductively coupled plasma atomic emission spectroscopy (ICP-AES) and surpasses the latter for refractory elements, such as Si because of incomplete solubilization during the digestion process (Gazulla et al., 2019; Reidinger et al., 2012). The XRF instrument relies on subjecting a sample with a beam of X-rays generated by an X-ray tube which causes X-ray fluorescence to occur in the sample (Jenkins, 1999). These excited fluorescent X-rays are then recorded and analysed to calculate the relative concentrations of elements present in the sample. X-ray fluorescence spectroscopy has low detection limits (<1 mg kg^−1^) for multiple elements of the Periodic Table, particularly the transition elements. However, the lower the atomic number of the element, the lower X-ray cross-section (efficiency of X-ray fluorescence generation) and the lower the energy of the fluorescent X-rays. This translates to a lower signal and attenuation (absorbtion) of the emitted fluorescent X-ray in the airpath between sample and XRF detector. This makes light element analysis with XRF extremely challenging (Marguí et al., 2022). In most desktop XRF instrumentation either vacuum or helium analysis conditions are used to minimize the effect of the latter.

While conventional XRF uses polychromatic excitation, the recently developed monochromatic XRF instrument used in this study (Z-Spec Inc. E-lite) has doubly curved crystals to produce a monochromatic excitation source (Chen and Wittry, 1998). This results in a dramatically lower background with a signal to noise ratio >1:1000 better than polychromatic XRF (Chen et al., 2008; Nadkarni et al., 2011). Practically, this translates to much lower limits of the detection, improved spectral deconvolution, and ease of quantification using a Fundamental Parameters approach (Kawai et al., 2019). The Z-Spec E-lite instrument incorporates monochromatic X-ray excitation at 4.5 keV (Ti-anode) with polarization and a silicon drift detector with ultra-thin carbon window to maximize low-energy detection (Z-spec). In a cellulose matrix (wheat flour) Z-Spec reports limits of detection (LODs) of 3 mg kg^−1^ for Si, 0.5 mg kg^−1^ for S, 0.4 mg kg^−1^ for Cl, 0.3 mg kg^−1^ for K, 0.3 mg kg^−1^ for Ca, and 20 mg kg^−1^ for Al. However, thus far the methodology had not been extended to Na yet, given the very XRF low energy (1.040 keV) making it perhaps the most challenging element to analyse using this method as it is strongly affected by sample and air attenuation of escaping fluorescent X-rays. Yet, it is the most crucial element to analyse in NaCl stress studies in plants.

In this study we developed a new method to accurately quantify light elements, including the toxic and beneficial ions Na and K in plant tissues (roots and shoots) using monochromatic XRF technology. In particular, we aimed to test the veracity of this technology for light element analysis (Na and K) in a range of different samples and plant species; to develop methods for sample preparation and mounting; and to evaluate its accuracy precision and repeatability. An additional objective was to test the feasibility of analysing very small sample amounts (<10 mg) generated from experiments involving young *Arabidopsis thaliana* plants with low biomass.

## RESULTS

### Development of the protocol

Detection of light elements heavily relies on extraction-based methods (typically involving acid digestion) requiring 5 mg or more of dried plant material followed by the most sensitive method available, ICP-MS. However, assuming a prevailing Na concentration of 2500 ppm (mg/kg) in the plant sample and 5 mg of sample material digested into 10 mL of solution, this translates to a analyte concentration of just 1.25 mg/L Na. When studying the early developmental stages to determine the perception of different salt stress stimuli, obtaining the required amount of material for a successful extraction is challenging. Therefore, methods which are non-extraction based and can utilize smaller amounts of dried plant material are instrumental for such studies. To address this, we pursued the use of monochromatic XRF to analyse the composition of plant material with minimal amounts (<10 mg).

We first tested the repeatability of this method by using NIST standards 1547 (Peach Leaves), 1570a (Spinach Leaves) and 1515 (Apple Leaves) representing different levels of Na and K. The amounts of 1mg and 10 mg in embedded form, where samples are packed between films (Table 1) and 250 mg (Table 2) packed using the plunger method, which compacts the sample into a sample cups for XRF analysis (see Figure 1). Ten measurements were performed for each sample with 60 seconds (s) measurement time. The results showed Na and K levels can be robustly detected even as low as in 1 mg, but the standard error was reduced in higher amounts of samples.

**Figure 1:**
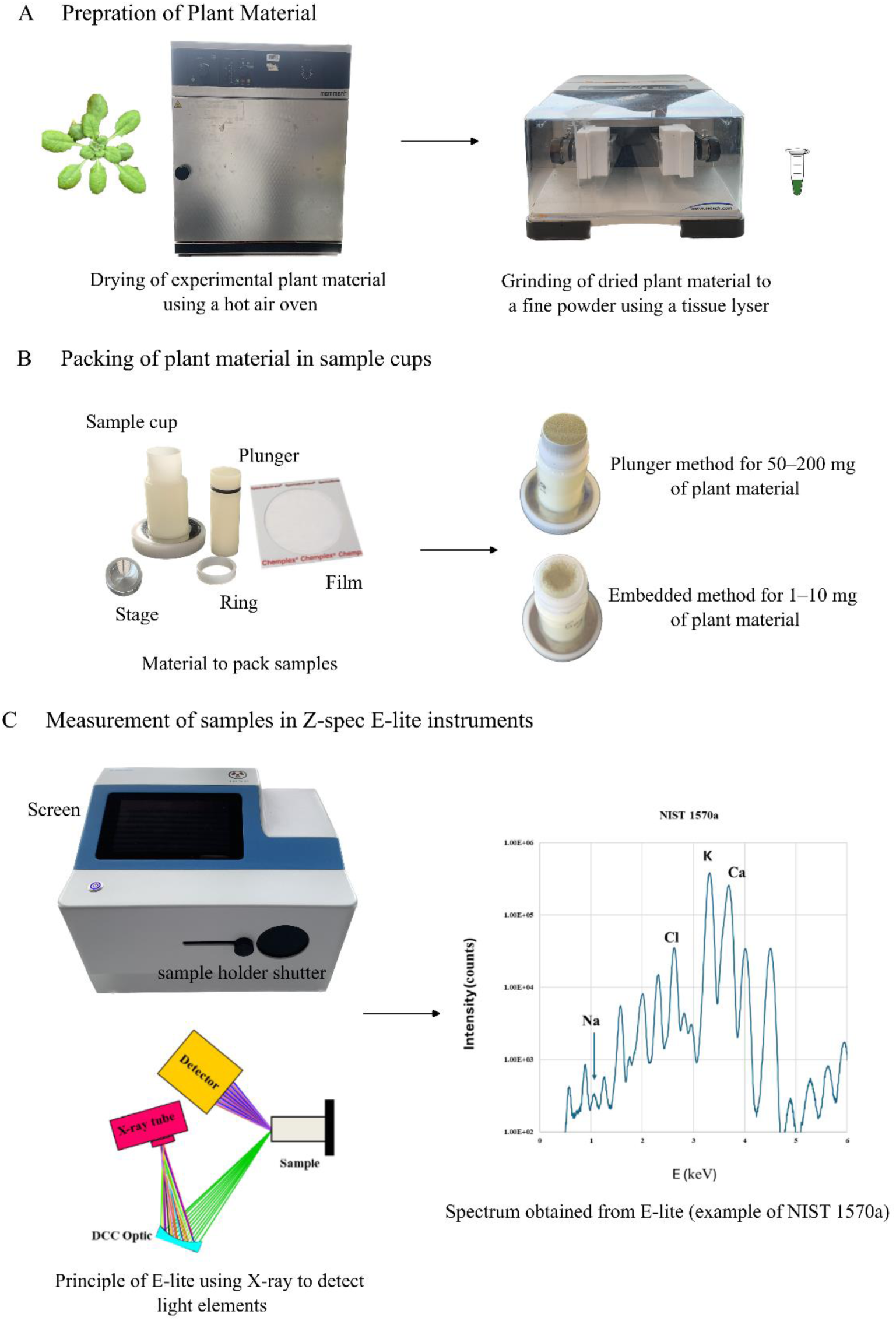
Schematic representation of the experimental procedure utilized for detecting low-energy elements using the Z-Spec E-lite instrument. The flowchart outlines the key steps involved in the analysis; A) including sample preparation; B) different material used for packing of dried plant sample; C) E-lite instrument, X-ray excitation, and the resulting data acquisition. The output spectrum is derived from the certified reference material NIST 1570a, highlighting the fluorescence peaks of Na, Cl, K and Ca.

**Table 1.**
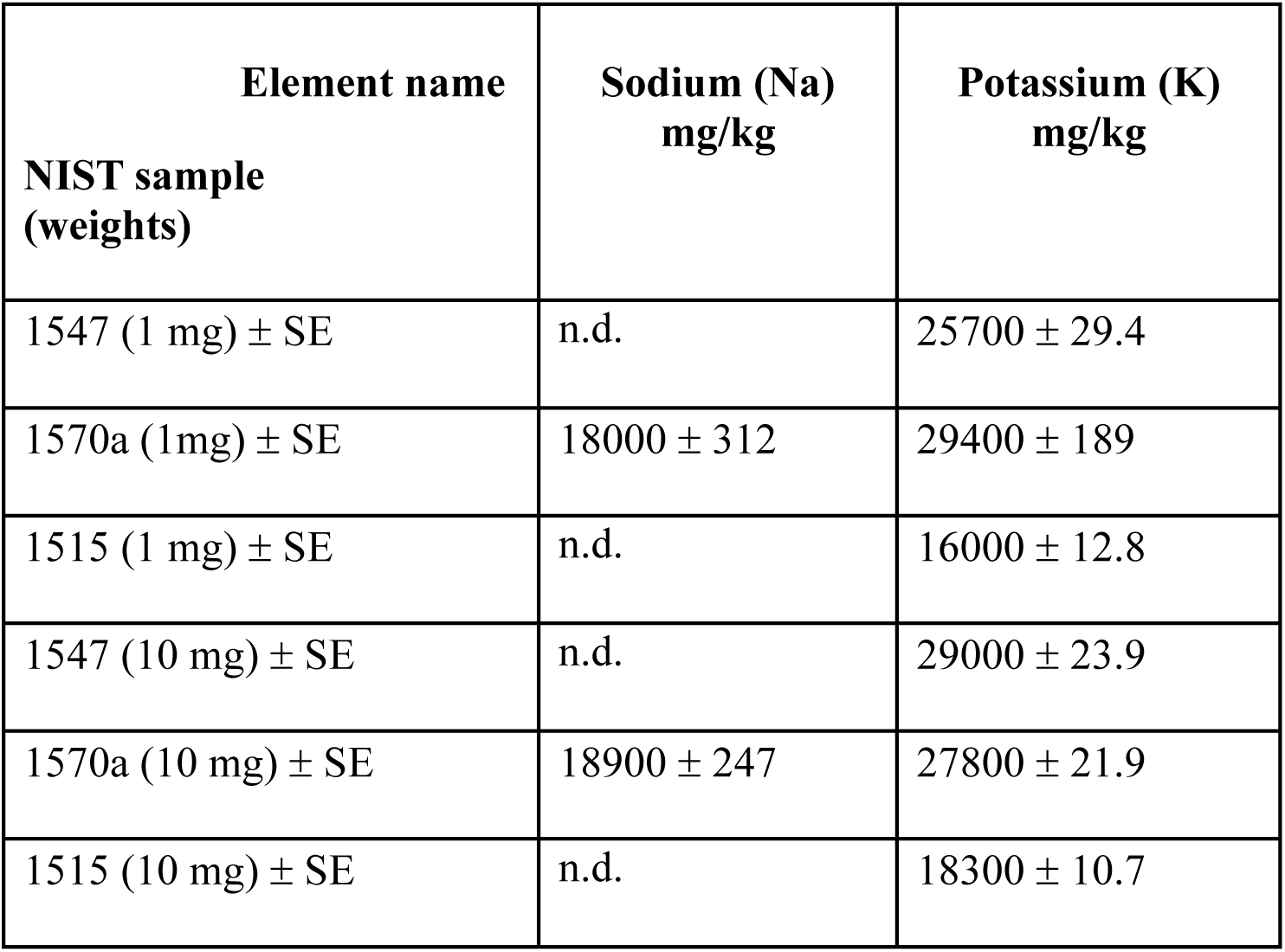
Embedded method for testing repeatability of XRF E-lite instrument using NIST standards 1547, 1570a and 1515, each sample was measured 10 times for 60 s and average of 10 samples is represented in table with standard error. The certified values of NIST 1547 for Na is 23.8 ± 1.6 and for K is 24330 ± 380; NIST 1570a for Na is and 18210 ± 230 and for K is 29000 ± 260; NIST 1515 for Na is 0 and for K is 16080 ± 210. n.d. is not detected.

| Element name<br>NIST sample<br>(weights) | Sodium (Na)<br>mg/kg | Potassium (K)<br>mg/kg |
| --- | --- | --- |
| 1547 (1 mg) ± SE | n.d. | $25700 \pm 29.4$ |
| 1570a (1mg) ± SE | $18000 \pm 312$ | $29400 \pm 189$ |
| 1515 (1 mg) ± SE | n.d. | $16000 \pm 12.8$ |
| 1547 (10 mg) ± SE | n.d. | $29000 \pm 23.9$ |
| 1570a (10 mg) ± SE | $18900 \pm 247$ | $27800 \pm 21.9$ |
| 1515 (10 mg) ± SE | n.d. | $18300 \pm 10.7$ |

**Table 2.**
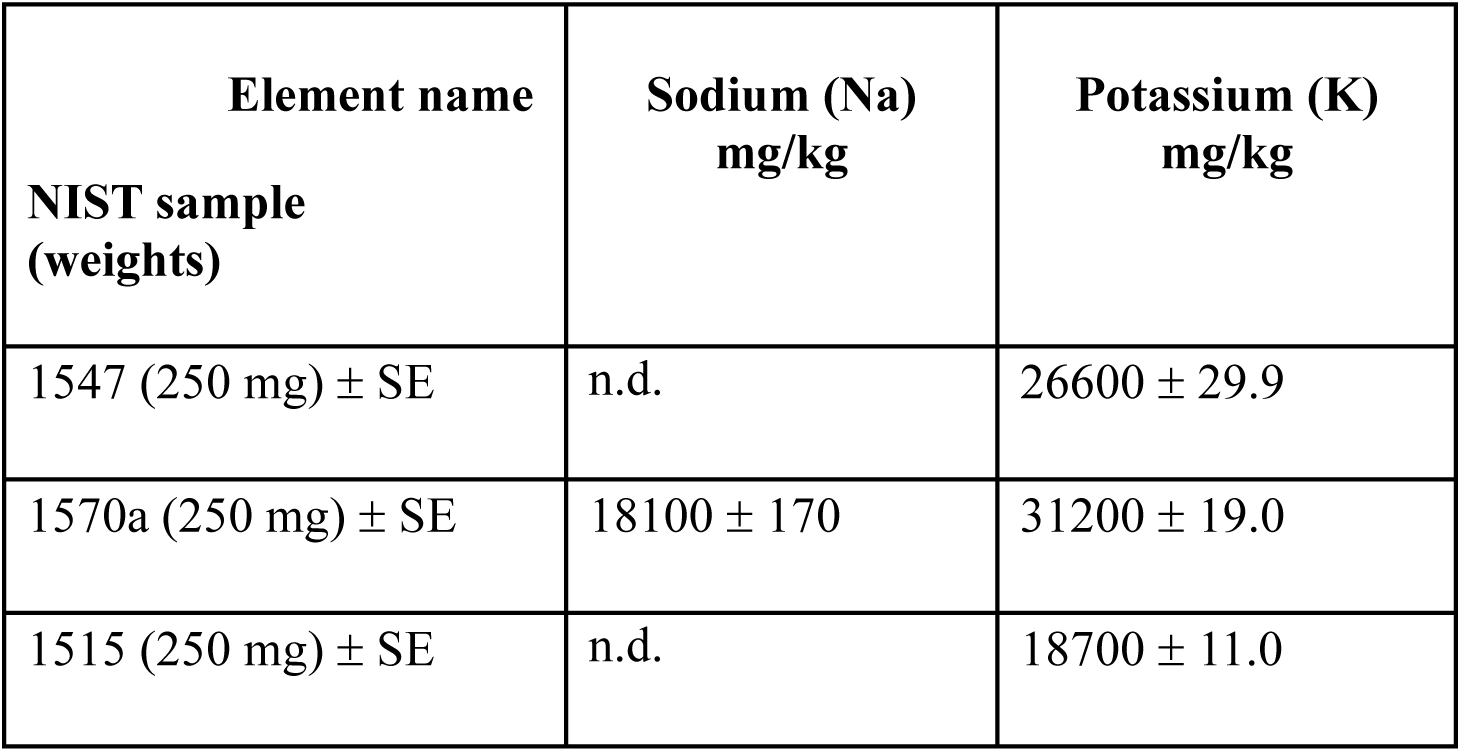
Plunger method with 250 mg of NIST standards 1547, 1570a and 1515 was performed to test repeatability of the E-lite instrument. Each sample was measured 10 times for 60 s and average of 10 samples is represented in table with standard error. The certified values of NIST 1547 for Na is 23.8 ± 1.6 and for K is 24330 ± 380; NIST 1570a for Na is and 18210 ± 230 and for K is 29000 ± 260; NIST 1515 for Na is 0 and for K is 16080 ± 210. n.d. is not detected.

Then we tested the amount of biological material required to measure Na and K precisely. We developed and tested two methods because different experimental conditions and biological samples often require varying amounts of material. The plunger method allows for larger sample sizes (5–200 mg) to accommodate more abundant tissues (Figure 2A), while the embedded method (1–50 mg) enables precise measurements with minimal material (Figure 2B), making it ideal for early developmental stages or scarce samples. The results showed for plunger method above 25 mg material was enough to reliably detect Na and K in *Arabidopsis thaliana* shoot samples. Although, 25 mg showed higher variance when compared to 50, 100 or 200 mg. For the embedded method 1–10 mg showed the best outcome. However, the 1 mg sample had higher variance when compared to 5 or 10 mg.

**Figure 2:**
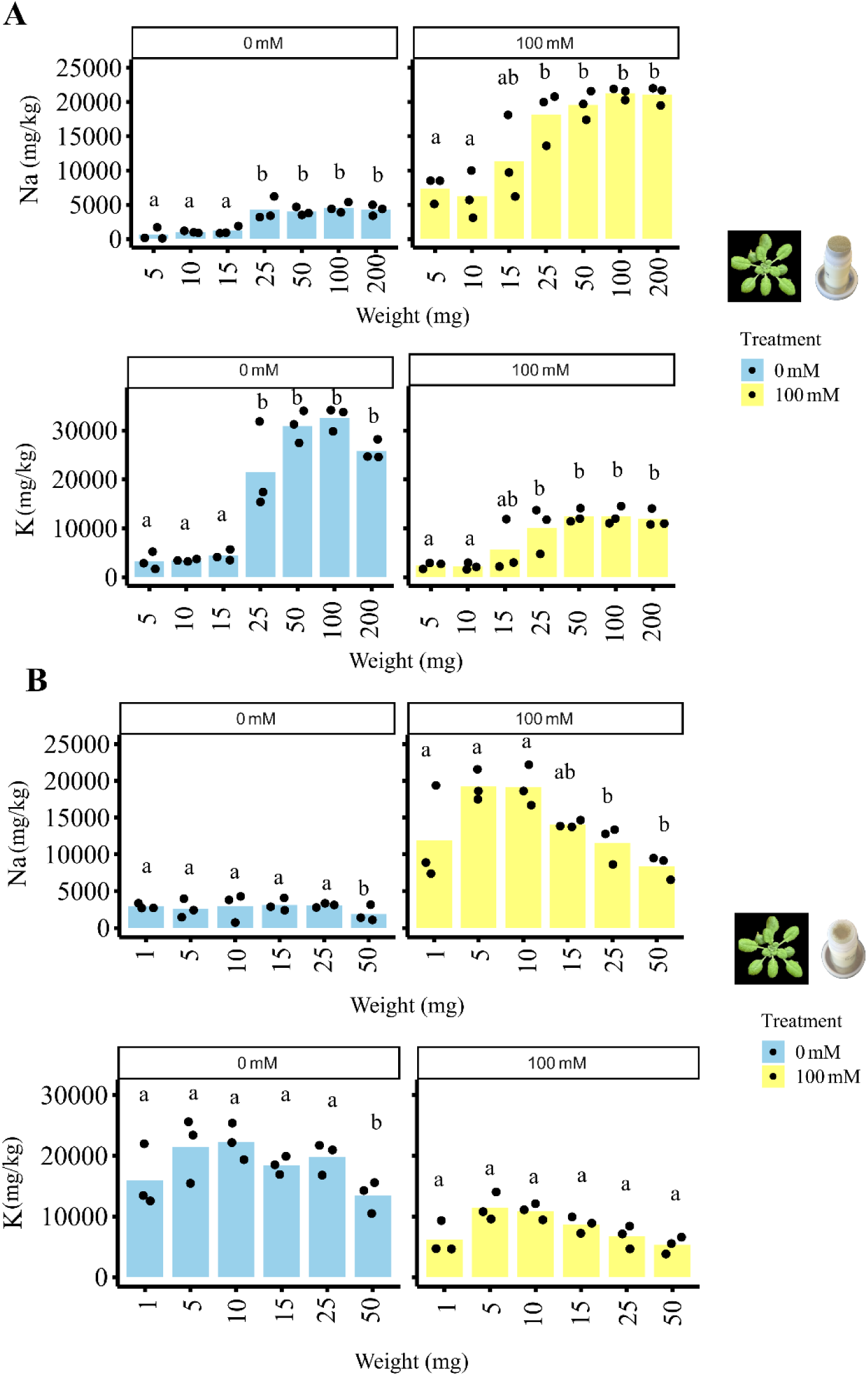
Amount of plant material required for Na and K analysis using two methods; A) plunger method (with plant material 5–200 mg) and; B) embedded method (with plant material 1–50 mg). The levels of sodium (Na) and potassium (K) are presented in mg/kg in *Arabidopsis thaliana* rosettes grown in two different conditions: control soil (blue bars) and plants watered with 100 mM NaCl (yellow bars). The varying amounts of plant material required for each method are compared in three independent replicates (represented by each dot). Data not sharing the same letter are significantly different.

Time of measurement was also tested for *Arabidopsis thaliana* and *Oryza sativa* shoot samples (Figure 3). Test time of 30, 60, 300 and 600 s was tested for the plunger and the embedded method. Our data showed no significant differences in the tested times for plunger method in soil grown *Arabidopsis thaliana* grown under control conditions or treated with 100 mM NaCl (shoot samples; Figure 3A). Similarly to the plunger method, we did not observe any differences in test time for the embedded method (sample preparation with 5 mg plant material) for soil grown *Arabidopsis thaliana* under control conditions as well as treated with 100 mM NaCl (shoot samples; Figure 3B) and hydroponics grown *Oryza sativa* sunder control conditions and supplemented with 50 mM NaCl (shoot samples; Figure 3C). Thus, our results indicate that neither the plunger nor the embedded method is sensitive to the duration of measurement within the tested time ranges for both *Arabidopsis thaliana* and *Oryza sativa* shoot samples. Suggesting that measurement time can be chosen based on practical considerations without compromising the accuracy of Na and K detection.

**Figure 3:**
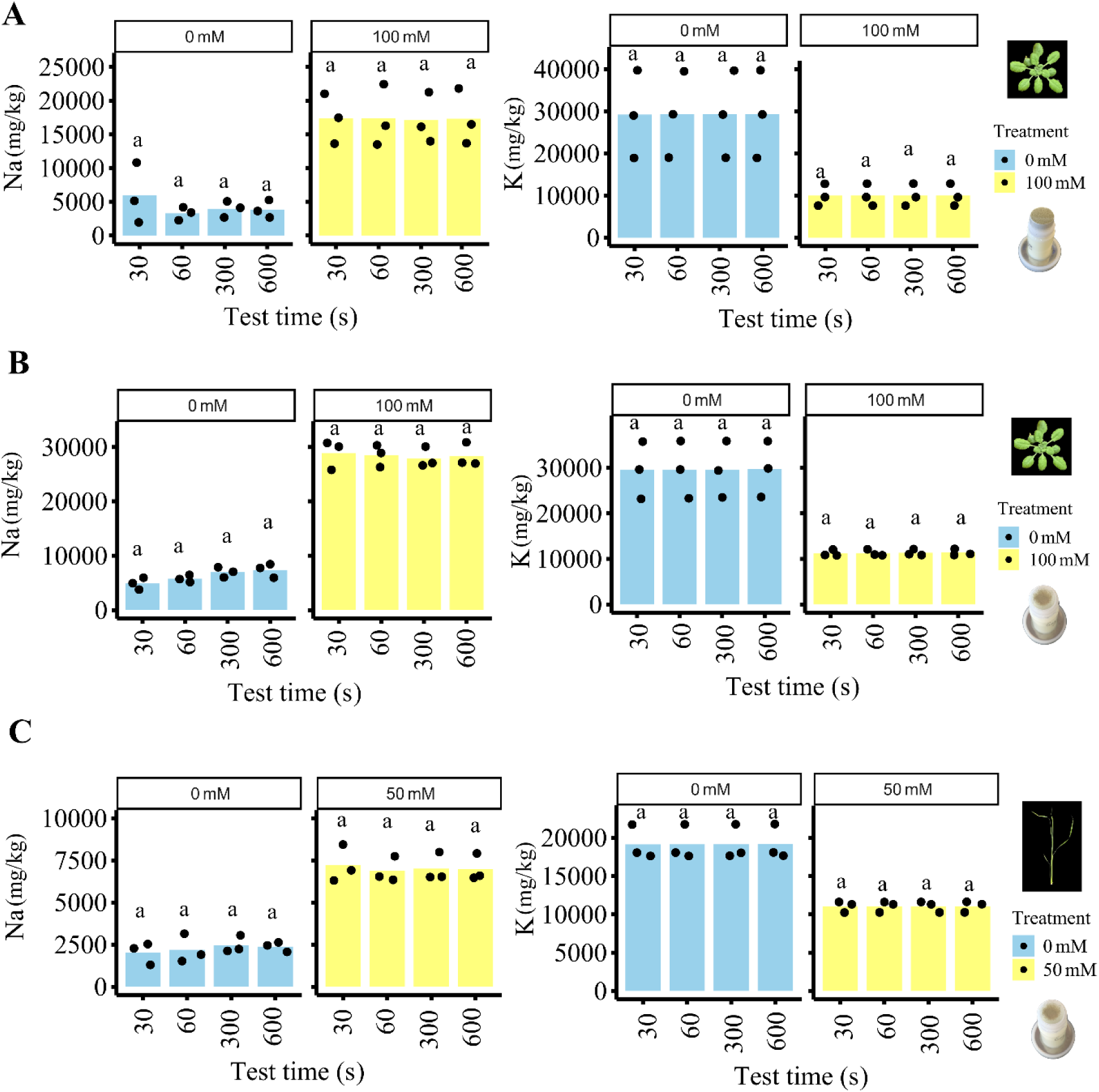
Time of measurement for sodium (Na) and potassium (K) analysis using two methods; A) plunger method (200 mg plant material) and; B) embedded method (5 mg plant material). The graph illustrates the duration tested for each measurement technique (30–600 s). Measurements are compared in *Arabidopsis thaliana* shoots grown under two conditions: control soil (blue bars) and plants treated with 100 mM NaCl (yellow bars). The different time of measurement for each method are compared in three independent replicates (represented by each dot). Data not sharing the same letter are significantly different.

Next, we compared data on Na and K obtained from XRF E-lite instrument to data obtained from a well-developed ion analysis method using ICP-MS (Figure 4). The dataset included *Arabidopsis thaliana* shoots grown in soil supplemented with 0, 75, 100, 125, or 150 mM NaCl solution and *Oryza sativa* seedlings (shoots and roots) grown on hydroponics solution with 0, 25 or 50 mM NaCl. Correlation plots were generated on data from *Arabidopsis thaliana* XRF measurements using the plunger method with 200 mg and 60 s measurement and corresponding ICP-MS data from the same samples (Figure 4A), XRF measurements using the embedded method (5 mg and 60 s measurement time) with the corresponding ICP-MS data from selected samples (Figure 4B) and *Oryza sativa* samples using the embedded method (5 mg) with the corresponding ICP-MS data from same samples (Figure 4C). All the data showed significant correlations with r values ranging from 0.86 to 0.92, raw data depicted in Supplemental Table 1.

**Figure 4:**
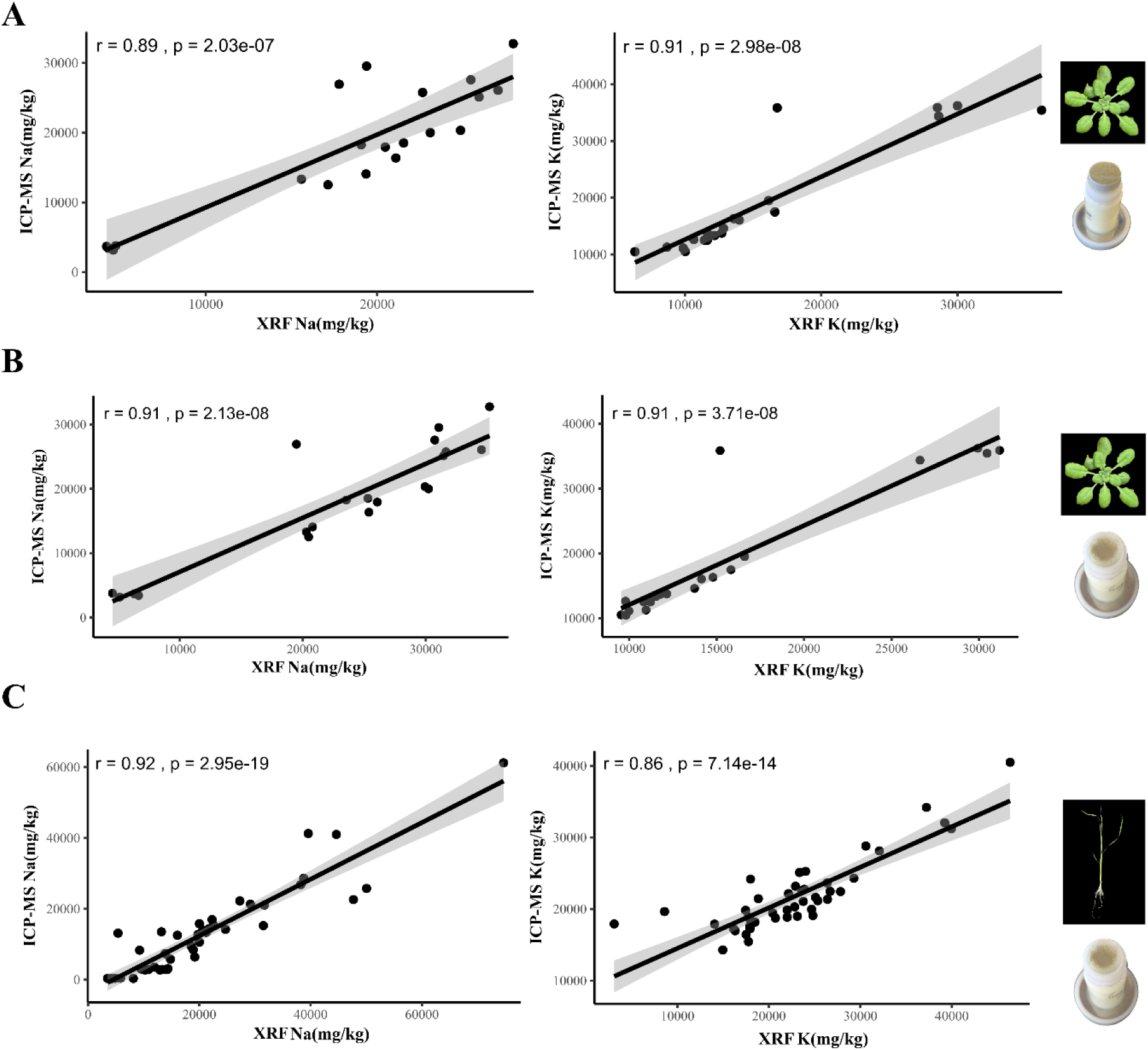
Comparison of measurement techniques for sodium (Na) and potassium (K) analysis using the; A) plunger method and; B and C) embedded method using monochromatic X-ray fluorescence (XRF) with ICP-MS. A) Data points represent measurements taken from *Arabidopsis thaliana* shoots treated with different NaCl concentrations using XRF plunger method (200 mg and 60 s) and ICP-MS (10–17 mg). B) Data points represent measurements taken from *Arabidopsis thaliana* shoots treated with different NaCl concentrations using XRF embedded method (5 mg and 60 s) and ICP-MS (10–17 mg). C) Data points represent measurements taken from shoots and roots of two *Oryza sativa* varieties treated with different NaCl concentration using XRF embedded (5 mg and 60 s) and ICP-MS (3–10 mg) under varying salt conditions. The graph illustrates the correlation between the results obtained from XRF and those from inductively coupled plasma mass spectrometry (ICP-MS) for both methods. The different treatments are compared in three independent replicates (represented by each dot), individual data point represents. Pearson correlation coefficients (r) are indicated, demonstrating the strength and direction of the linear relationship between the datasets for each method.

### Proof of concept testing of monochromatic XRF analysis

Finally, we performed a proof-of-concept experiment using roots of *Arabidopsis thaliana* Col-0 seedlings, which were grown on agar plates for 7 days before being treated with 125 mM NaCl for 6 hours. This treatment allowed us to assess the effects of salt on root ion levels by comparing the sodium (Na) and potassium (K) levels to those of corresponding controls, including samples shifted to NaCl plates and harvested immediately; 0 hours samples and samples that were not exposed to NaCl for the 6-hour period (Figure 5A).

**Figure 5:**
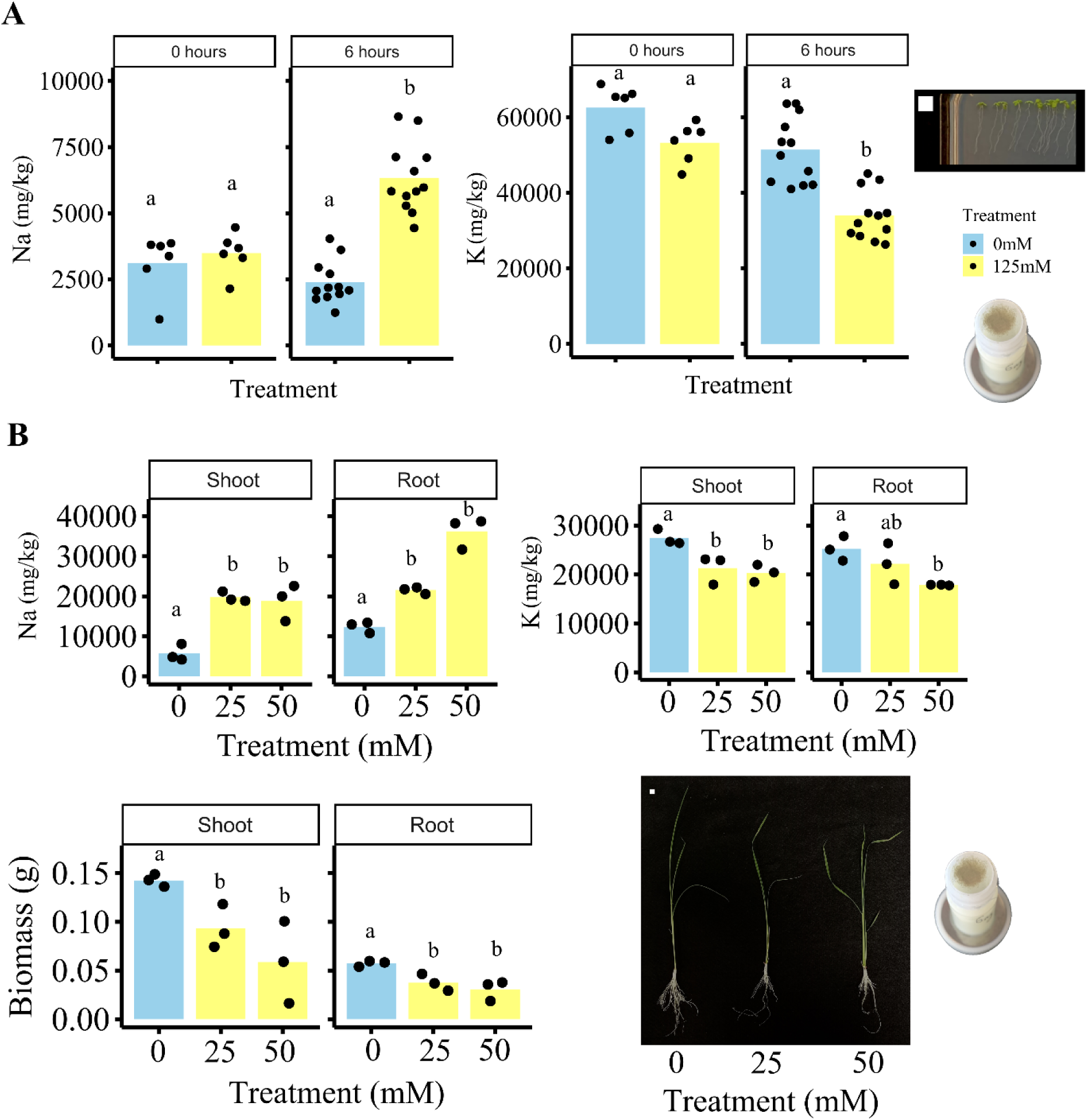
Proof-of-concept results demonstrating effects of salinity on sodium (Na) and potassium (K) levels using embedded method with 5 mg and 60 s measurement. A) *Arabidopsis thaliana* grown on agar plates treated with NaCl for 6 hours, highlighting the early uptake of salt in root tissue. The image shows a representative agar plate with plant samples, 1×1 cm white square. B) Effects of treatment with two different salt concentrations (25 and 50 mM) for 14 days on *Oryza sativa* variety Mudgo, illustrating Na and K levels in both shoot and root tissues. The relationship between NaCl concentration and root biomass is depicted, underscoring the impact of salinity stress on growth. Blue bars represent control conditions and yellow bars represent different salt treatments, each data point is an independent replicate, with letters above the bars indicating statistical significance; groups not sharing the same letter are significantly different from each other.

The data obtained from the root tissues indicated a significant increase in Na concentration and a corresponding decrease in K levels following the 6-hour NaCl treatment. This shift in ion balance suggests that the plant is responding to the saline environment by accumulating Na at this early timepoint. Such alterations in early ion homeostasis are critical to understanding the mechanisms plants use to cope with salt stress.

To further explore the effects of salinity on plant performance, we conducted a salinity experiment with the Mudgo variety of *Oryza sativa*. The plants were grown in a hydroponic system, with some groups receiving a control nutrient solution and others supplemented with either 25 mM or 50 mM NaCl for 14 days (Figure 5B). The results from this experiment showed a significant increase in Na levels in both NaCl treatments. Simultaneously, we observed a decrease in K concentrations, which was accompanied by a reduction in both shoot and root dry biomass.

Overall, our proof-of-concept experiments offer valuable insights into the use of *Arabidopsis thaliana* roots at a very early developmental stage where plant material is limited, as well as showing correlation between mild salinity responses and dry biomass in both *Oryza sativa* shoots and roots.

## DISCUSSION

The results of this study demonstrate that the monochromatic XRF method is highly effective for quantifying Na and K concentrations in small amount of plant samples, providing a reliable, non-extraction alternative to traditional extraction-based techniques like ICP-MS. X-ray florescence analysis has been shown to be a powerful tool for the analysis of various elements in biological and environmental samples due to its high precision, non-destructive nature, and ability to measure low concentrations of elements (Castillo-Michel et al., 2017; Maliki et al., 2017; Marguí et al., 2005). X-ray florescence analysis has been successfully used in the past for performing multi-elemental analysis and has shown promising outcomes with both Na and K, with larger plant samples, approximately up to 2 grams of plant material (Marguí et al., 2005). In the current method, the ability to analyse as little as 1–10 mg of plant material is primarily attributed to advancements in (polarised) monochromatic XRF technology itself, which offers enhanced sensitivity and precision compared to polychromatic XRF that is commonly used. Instrumental advancements incorporated in the E-lite analyser include very short sample to detector geometry, newest generation of low-energy optimised silicon drift detector with graphene window, and ultra-low noise electronics. Moreover, the use of an innovative method for sample mounting using a thin sample layer in a film sandwich approach ensures a very low scattering background further improving the signal to noise ratio. This was further improved by utilizing a background subtraction and Compton Scatter correction in the Fundamental Parameters calculation for each sample.

Another important aspect of XRF analysis is sample preparation, including the type, amount, and thickness of the sample. Various preparation methods can be used, such as pressed pellets and loose powders, with smaller particle size to enhance analytical accuracy but all requiring larger amounts of samples (Marguí et al., 2022). In contrast, we present a method that requires only 1–10 mg of plant material, embedded between two films. This approach addresses previous limitations by enabling precise measurements with minimal sample sizes, thus facilitating the analysis of scarce biological materials.

The significant advancement of our method lies in both the sophisticated monochromatic XRF technology and the innovative sample preparation technique the allows for the analysis of minimal sample sizes. Significant correlations between XRF and ICP-MS data for samples from both *Arabidopsis thaliana* and *Oryza sativa* experiments validate the accuracy of this method across different sample types and treatment conditions. The method we present here has the ability to measure Na and K in as little as 1 mg of plant material opening up new possibilities for high-throughput analysis, particularly in experiments involving early plant developmental stages or limited biological material.

### Applications of the method

The study focuses on light elements, mainly Na and K, which are analysed in roots and shoots of *Arabidopsis thaliana* and *Oryza sativa*. Yet, the method has wider potential applications for a variety of tissues like flowers to seeds. In addition, soil and liquid samples can also be tested and analysed to assess nutrient use efficiency comprehensively. Moreover, the method’s ability to provide simultaneous multi-elemental analysis makes it particularly valuable for understanding nutrient uptake dynamics over time, especially when analysing liquid samples. Validation of results obtained through XRF has been shown to correlate well with established techniques such as inductively coupled plasma mass spectrometry (ICP-MS), underscoring the robustness and reliability of this approach (Singh et al., 2021). The method offers a versatile approach, applicable to different types of plant tissues for determining the uptake, distribution, and accumulation of essential elements without the need of extractions and can be determined quantities as low as 1 mg (Figure 2). Small and large quantities of plant materials can be processed, providing flexibility depending on the experimental needs and sample availability. Liquid samples are particularly valuable for understanding nutrient uptake dynamics and efficiency over time. For validation and comparison, the results obtained using this method have been correlated with data from ICP-MS (Figures 4). This comparative analysis highlights the robustness of our approach against established analytical techniques.

In this article, we focus on using this method to study plant leaves and roots under NaCl stress conditions. This allows us to assess toxicity, uptake, and accumulation patterns across different plant species.

### Limitations of the method

While this method has demonstrated versatility and reliability, there are some inherent limitations that must be addressed for optimal performance. One notable limitation is the vulnerability of the detector to dust contamination that can affect the uniformity of results, necessitating regular cleaning to maintain accuracy and to prevent contamination. This requirement can add to the maintenance workload and can influence the consistency of results if not properly managed. Additionally, the method requires the use of highly uniform and finely ground dry powder (finer than 0.1 mm particles) from the sample material, as particle size has been shown to have a significant effect on the accuracy of the measurements using XRF analysis (Amar et al., 2022; Beckhoff et al., 2006). Achieving this level of uniformity can be challenging, especially for different types of plant tissues such as roots, shoots, leaves, and seeds. Any inconsistency in particle size may lead to variability in the results, thus emphasizing the importance of meticulous sample preparation.

## CONCLUSIONS

The monochromatic XRF method approach reported here is a new and robust method for the routine analysis of light elements such as Na and K in plant tissues. It offers significant advantages, including the ability to analyse small sample sizes with minimal preparation and no hazardous chemicals. This makes it particularly attractive for applications ranging from breeding programs to ecological research and this study successfully developed and validated a monochromatic XRF-based method for the rapid and accurate quantification of Na and K across various plant tissues. By eliminating the need for chemical digestion and allowing for the analysis of very small samples, the method is highly suitable for investigating plant responses to stress, nutrient management, and salinity tolerance, with potential applications in agriculture, plant breeding, and environmental science. Overall, despite some challenges in sample preparation, this technique represents a promising tool for future research in plant light elemental analysis and will be highly suited for investigating other elements of importance in physiology such as silicon (Si), aluminium (Al), phosphorus (P) and sulfur (S).

## EXPERIMENTAL PROCEDURE

Materials: Sample cups, rings, 4.0 μm film, dried plant material (depending on the amount of the plant material select one of the following methods).

Equipment: E-lite XRF (Z-Spec Inc., https://zspecinc.com/hdxrf), utilizing monochromatic X-ray excitation at 4.5 keV (Ti-anode) with polarized radiation, and a silicon drift detector featuring an ultra-thin carbon window to maximize low-energy detection (Figure 1A-C).

### Preparation of sample cups (50–200 mg sample material) Plunger method

- Place a layer of film at the top of the sample cup and secure with a ring (sample cup, film and ring shown in Figure 1B).
- Cut off the excess film with a sharp scalpel.
- Place the sample cup upside down on the stage (shown in Figure 1B).
- Add dried ground sample inside the sample cup to a 3-5 mm thickness or approximately 200 mg.
- Press the plunger inside the sample cup (shown in Figure 1B) to pack the sample. Note: Avoid any kind of non-uniform packing as that will lead to errors in measurements.
- Clean the sample cups for any residues to avoid spill overs inside instrument.

### Preparation of sample cups (1–15 mg) Embedded method

- Place a layer of film at the top of the sample cup and secure with a ring (sample cup, film and ring shown in Figure 1B).
- Cut off the excess film with a sharp scalpel.
- Make sure to remove any bits and pieces of excess film, to avoid spills inside the machine.
- Put the finely ground sample on top of the film.
- Place a second layer of film at the top of the sample and use a second ring to embed the sample in-between the two films. Cut off the excess film. Note: Avoid any kind of non-uniform packing as that will lead to errors in measurements.
- Clean the sample cups for any sample residues to avoid any spill overs inside the machine.

### XRF Elite measurement

- Switch on the machine by pressing the ‘switch on’ button for 3 s and wait for 30-60 s for initialisation.
- Set-up the parameters of experiment under ‘Set-up’ on the initial screen (see Figure 1).
- Standard parameters are: 60 s test time, 30 s interval time.
- Open the sample cup holder shutter (see Figure 1C) and remove the stopper from front of the detector.

Caution: whenever opening the shutter make sure the X-ray light indication on top of the machine is always off.

- Clean the sample area from the dust particles. Note: It is to avoid any accumulation of dust on the detector as it will heavily influence its efficiency, but do not clean or touch the detector itself. Make sure the sample cup to be measured is clean.
- Place the sample cup into the cup holder in instrument and close the lid. Note: Be gentle as the sample cup holder and shutter are magnetic.
- Press the start tab on the screen for starting the measurement Caution: Make sure the X-ray light is off, and the status of the measurement is ‘finished’ before opening the lid and inserting the next sample for the measurements.

### Retrieving and processing the data

- Once all the measurements are complete, export the data as follows: history/export/curve name=material selected while setting up the parameters of the experiment and export the csv file. Firmware can export up to 200 measurements in one single csv. File. If more measurements need to be exported, data can be extracted by getting single excel files and transforming them manually. That can be performed as follows: transfer/select the data you want/copy. Note: Avoid the move function to avoid unintentional loss of data.

## MATERIALS AND METHODS

### Arabidopsis thaliana soil experiment

The *Arabidopsis thaliana* Col-0 ecotype plant material was obtained from a salt tolerance assay performed on soil. Seeds were stratified at 4°C in the dark for 3 days and they were sown on soil in 7×7 cm pots (48 replicates/treatment). The experiment was performed in a climate chamber with 16 hours light, 8 hours dark, 20°C/18°C temperature, 60% humidity and light intensity of 130 µmol m^2^ s^−1^ PAR. The salt treatment was performed as in Bellinazzo et al., 2024 with some modifications. In this experiment, the pots were watered with tap water from the bottom, then 0.5 L of tap water containing the indicated salt concentration (0, 75, 100, 125 or 150 mM) together with the water left from the last watering step was added per tray of 24 pots from the bottom and thereafter they were watered with tap water and Hyponex solution (S Table 2). Whole rosettes were harvested after 21 days of treatment and dried at 60°C for 1 week before weighing. Twelve rosettes per treatment were pooled and ground together constituting a single sample.

### *Oryza sativa* hydroponics assay

*Oryza sativa* seeds (rice) from two varieties (Mudgo and M. Baltec) were obtained from the International Rice Research Institute (IRRI), Philippines. Seeds were dehulled and sterilized with 70% ethanol for 1 minute, followed by 30% commercial bleach solution for 25-30 minutes, then washed with MQ water 5 times. Germination was conducted using 0.3% gel rite (Duchefa Biochemie) with Yoshida nutrient solution (S Table 3; Yoshida et al., 1976) in modified PCR tubes, incubated in hydroponic boxes at 37°C for 24 hours and then at 28°C for 24 hours. Plants were grown in climate chamber at 28°C/25°C (day/night) with 75% humidity and 400µmol m^−2^ s^−1^ PAR light. Seedling were grown in a hydroponic setup using Yoshida nutrient solution (pH 5.2) for 14 days supplemented with NaCl (0, 25 and 50 mM) were used to test salinity response. The root (washed three times with MQ) and shoot were harvested and dried at 70°C for 3 days to determine dry weights.

### Arabidopsis thaliana plate assay

*Arabidopsis thaliana* Col-0 seeds were surfaced-sterilised using 70% Ethanol for 1 min followed by 10 min of sterilisation medium (30% of commercial bleach, 0.02% Triton X-100, Sigma-Aldrich). Then they were sown on ½ Murashige and Skoog (MS) medium including vitamins (Duchefa Biochemie), supplemented with 1.1 g/L MES (Duchefa Biochemie) and 1% Diashin Agar (Duchefa Biochemie), pH 5.8. After 3 days of stratification in dark at 4°C, plates with seed were then placed vertically at 90° in a growth chamber for 7 days in white light (100 µmol m^−2^ s^−1^ PAR, 16 hours photoperiod, 20°C temperature). At day 7, seedlings were transferred to ½ MS with or without 125 mM NaCl for indicated time period. Roots samples were collected and rinsed with milliQ water three times. Samples were then dried at 60°C for 1 week.

### Elemental analysis of plant samples

Dried root and shoot samples were then placed in a 2 mL Eppendorf tube or 12 mL tube with 5–10 metal beads (0.8mm). The plant material was then ground to a fine powder (<200 µm) in a tissue lyser (Retsch) and subsamples were inserted into custom XRF sample cups and covered with a layer of 4 µm polypropylene thin film (Chemplex Industries Inc.) for XRF analysis. The XRF analysis of the plant powdered material was performed using a E-lite instrument (Z-Spec Inc.). The instrument use monochromatic X-ray fluorescence excitation at 4.5 keV to analyse elements Z = 11 (Na) to Z= 20 (Ca). Samples were analysed for 30–600 s in plant (for samples prepared with plunger method) or thin sample (for samples prepared with embedded method) mode. Quality controls included NIST SRM 1547, 1570a, and 1515. Inductively coupled plasma mass spectrometer (ICP-MS) analysis was performed using samples ranging from 3–17 mg. Ion measurements were conducted at the Ionomics Facility, University of Nottigham, UK, following the methodology with minor modifications from Danku et al. (2013). Samples were digested by adding 1 mL of trace analysis grade (TAG) concentrated HNO₃ to each tube and predigesting overnight in a fume hood, followed by the addition of 0.5 mL of 30% H₂O₂ and digestion for 4 hours at 115°C on a block heater. The samples were diluted to 10 mL with Milli-Q water (18 MΩ). Multi-element analysis was carried out using ICP-MS (Thermo Fisher Scientific iCAP-Q, Bremen, Germany) with a flow rate of 1.2 mL min⁻¹ via an autosampler (Cetac ASX-520) and an ASXpress™ rapid uptake module, using a perfluoroalkoxy (PFA) Microflow PFA-ST nebulizer. Data were processed with Qtegra™ software, applying external cross-calibration between pulse-counting and analog detector modes. The system employed in-sample switching between two modes: (i) helium gas with kinetic energy discrimination (KED) to remove polyatomic interferences and (ii) hydrogen gas for selenium determination. Internal standards (Sc and Ge at 10 µg L⁻¹, Rh and Ir at 5 µg L⁻¹) were introduced via the ASXpress at an equal flow rate to correct for instrumental drift. Calibration included (i) a multi-element standard (Ag, Al, As, Ba, Be, Cd, Ca, Co, Cr, Cs, Cu, Fe, K, Li, Mg, Mn, Mo, Na, Ni, P, Pb, Rb, S, Se, Sr, Ti, Tl, U, V, Zn) at 0–100 µg L⁻¹ (Claritas-PPT grade CLMS-2, SPEX Certiprep, Metuchen, NJ, USA), (ii) a multi-element calibration solution (PlasmaCAL, SCP Science, France) with Ca, Mg, Na, and K at 0–30 mg L⁻¹, and (iii) an in-house prepared mixed phosphorus, boron, and sulfur standard (KH₂PO₄, K₂SO₄, H₃BO₃). Certified reference material NIST 1573a (Tomato Leaves) was digested alongside samples and blanks for quality control..

### Statistical analysis

Data analyses in this study were performed in R software (version 4.2.0, R Core Team, 2023). A *p-value* of less than 0.05 was considered statistically significant. The normality of the data for each treatment group was assessed using the Shapiro-Wilk test shapiro.test(). If the data followed a normal distribution, an ANOVA aov() was performed to compare the groups at a 95% confidence level. In cases where significant differences were detected, a post-hoc Tukey test TukeyHSD() was performed. However, if the data did not meet the normality assumption, the non-parametric Kruskal-Wallis test kruskal.test() was applied, followed by Dunn’s test dunnTest() for post-hoc comparisons. Pearson’s correlation analysis was executed using the cor() function. Plots were created with the R package ggplot2.

## AUTHORS CONTRIBUTIONS

PSK, AvdE, and CT: conceptualization; PSK, PK, SBA, and CKL: investigation; PSK, AvdE, CT, PK, SBA, CKL, and ZC: data interpretation and evaluation; PSK and AvdE: writing and data representation. All authors critically reviewed and approved the manuscript.

## ACKNOWLEDGEMENTS

We would like to thank Nafiatul Umah from Laboratory of Plant Physiology, Wageningen University and Research, Netherlands for her help in conducting the *Oryza sativa* hydroponic experiments, Joram Dongus from Laboratory of Plant Physiology, Wageningen University and Research, Netherlands for his guidance during the *Arabidopsis thaliana* soil experiments. We also thank Liz Bailey and Lolita Wilson from School of Biosciences, University of Nottingham, United Kingdom for ion measurements and Mariana Silva Artur from Laboratory of Plant Physiology, Wageningen University and Research, Netherlands for her insightful feedback on the manuscript.

